# Group size influences individual metabolic traits in a social fish

**DOI:** 10.1101/2021.04.20.440644

**Authors:** Emmanuelle Chrétien, Daniel Boisclair, Steven J. Cooke, Shaun S. Killen

## Abstract

1. Group living is widespread among animal species and yields both costs and benefits. Presence of conspecifics can restrict or enhance the expression of individual behaviour, and the recent social environment is thought to affect behavioural responses in later contexts, even when individuals are alone. However, little is known about how social dynamics influence the expression of individual physiological traits, including metabolic rates.
2. There is some evidence that shoaling can reduce fish metabolic rates, but habitat conditions such as shelter availability may generate density-dependent influences on individual metabolic rates.
3. We investigated how social group size and availability of shelter influence Eurasian minnow *Phoxinus phoxinus* metabolic rates estimated by respirometry in the presence or absence of plant shelter. Respirometry trials were conducted before and after we housed fish for three weeks in a social treatment consisting in a specific group size (n= 4 or 8) and shelter availability (presence or absence of plant shelter in the holding tank).
4. Minimum day-time and night-time metabolic rates estimated while in presence of plant shelter were lower than when estimated in absence of plant shelter, both before and after individuals were housed in their social group size and shelter availability treatment. Standard metabolic rate was higher for fish held in groups of four as compared to fish held in groups of eight while maximum metabolic rate showed no difference. Shelter availability during the social treatments did not influence standard or maximum metabolic rates.
5. Our results suggest that group size may directly influence energy demands of individuals, highlighting the importance of understanding the role of social dynamics on variations in physiological traits associated with energy expenditure.

## Introduction

An animal social group is any set of individuals that remain together in space and time (Krause & Ruxton, 2002). Group living can provide a number of benefits, such as reduced predation risk, improved foraging, increased mate choice, and reduced energetic cost of movement or thermoregulation (Evans et al., 2016; Jolles et al., 2020; Krause & Ruxton, 2002). Conversely, group living can be associated with increased conspicuousness or attack rates from predators, reduced individual growth if food resources are limited, and increased parasite or disease burden (Altizer et al., 2003; Guénard et al., 2012; Hoare et al., 2004). Social structures emerge in groups from variability in individual behaviour and interactions among groupmates. Some behavioural responses are influenced by the number of groupmates present (Krause & Ruxton, 2002). For example, group size has been negatively correlated with foraging in novel contexts (Day et al., 2001) but positively correlated with exploration (Ward, 2012). Presence of conspecifics can restrict or enhance the expression of individual behaviour through processes like conformity or facilitation (Jolles et al., 2016; Ward, 2012; Ward & Webster, 2016). Consequently, individuals may express a different suite of behaviours and different degrees of their full behavioural capacity while in group compared to when they are alone (Jolles et al. 2020). Further, there is some evidence that the recent social environment can affect behavioural responses in later contexts, even when individuals are alone (Jolles et al. 2016). This suggests that the social environment could modulate an individual’s behavioural expression or capacity, yet the ways in which the phenotype of individual animals interact with their social environment remains largely unknown, including how social dynamics affect individual physiological traits.

The interplay between the social environment and individual physiological traits may be especially complex due to the effects of social dynamics on individuals stress, energy intake, and energy use (Webster & Ward, 2011). For instance, standard metabolic rate (SMR), the minimum rate of energy use needed to sustain life at a given temperature in an ectotherm (Burton et al., 2011; Chabot et al., 2016), generally correlates with dominance, aggression, and tendency to take risks among individuals (Arnold et al., 2021; Biro & Stamps, 2010; Metcalfe et al., 2016; Redpath et al. 2010). However, there is also evidence that individual stress can influence SMR over various temporal scales. In brown trout *Salmo trutta*, holding in pairs led to an increase in SMR of subordinate individuals, probably due to social stress, while SMR of dominant individuals did not change (Sloman et al., 2000). This is an example of how dominance can modulate relationships between metabolism and behaviour (Killen et al., 2013), though whether such effects occur in larger or more complex social systems than dyads requires further investigation. There is evidence, however, that shoaling can reduce SMR in fish through “calming effects” (Nadler et al., 2016). Like SMR, maximum metabolic rate (MMR) and aerobic scope (AS; the difference between MMR and SMR) can correlate with dominance (Killen et al., 2014), boldness, or aggression (Redpath et al. 2010). However, to our knowledge, there is no evidence to date that social stress can influence MMR or AS (Killen, Croft, et al., 2016), despite their potential to constrain energetically costly behaviours and other aerobically fueled activities (Metcalfe et al., 2016). In any case, SMR and MMR are often positively correlated (Auer et al. 2017; Killen, Glazier, et al., 2016; Norin & Clark, 2016) within and across species. As such, any effects of social dynamics on metabolic rates at rest may also affect aerobic capacity, or vice versa. The potential for social dynamics to influence either SMR or MMR could be reflected in AS, and thus influence the capacity to perform aerobically fueled activities. Yet, few studies have investigated how group living affects interactions between behavioural and physiological traits (Huang et al., 2020), aside from studies looking at effects of dominance in dyads (Sloman et al., 2000).

Habitat may further modulate interactions between individual traits and social dynamics (Jolles et al., 2020). Habitat conditions such as temperature or oxygen concentration influence metabolic rates, which in turn may affect performance among individuals within groups (Claireaux & Lefrançois, 2007; Fry, 1971; Horodysky et al., 2015; Huey, 1991). Conversely, social stress can reduce tolerance to thermal stress (LeBlanc et al., 2011) and hypoxia (Thomas & Gilmour, 2012). Other habitat conditions such as food and shelter availability may exert density-dependent influences on relationships between metabolism and behaviour. A number of studies have revealed that SMR or RMR estimated while in presence of shelter were reduced compared to when shelter was absent, probably due to decreased stress or reduction of alertness or vigilance when individuals are visually hidden (Chrétien et al., 2020; Finstad et al., 2004; Fischer, 2000; Millidine et al., 2006; Norin et al., 2018). However, little is known about the effects of long-term shelter availability on individual metabolic rates and interactions with an animal’s social environment. Increased competition for a limited resource, like availability of shelter, could strengthen social hierarchies and increase stress experienced by subordinates, and these effects could be greater in larger social groups. As such, group size and long-term shelter availability may have interacting effects that carry over and influence individual metabolic rates.

We investigated whether exposure to a given group size and shelter availability could influence metabolic rates of Eurasian minnows *Phoxinus phoxinus,* a small Cyprinid naturally living in social groups (Magurran, 1986). We held fish in groups of four or eight, in tanks with or without plant shelter. The combination of group size and shelter availability in holding tanks generated social treatments that differed in fish density and potential competition intensity for use of shelter. Respirometry trials were conducted before and after fish were housed for three weeks in these different social treatments, to measure metabolic rates 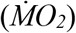 in presence or in absence of plant shelter. This design allowed us to get estimates of day-time and night-time 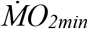 in presence or in absence of plant shelter, as the importance of being visually hidden by a shelter may vary with light intensity, as well as estimates of SMR, MMR, and AS. We hypothesized that the recent social environment would have metabolic costs that carry over, even when individuals are alone (Jolles et al., 2016), and be reflected in estimates of metabolic rates. Consequently, we predicted that presence of plant shelter during respirometry trials would lower day-time 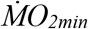, but that the magnitude of this effect would be smaller after the fish were held for three weeks in their social treatment (Killen et al., 2013). Given that minnows are social fish, we also predicted that SMR would vary with group size, due to the potential for social dynamics to modulate SMR (Sloman et al., 2000). We also predicted that fish held without access to plant shelter would have higher SMR, due to chronic effects of stress (Huey, 1991). The potential for group size and shelter availability to influence MMR is unclear. One the one hand, MMR is generally thought to be less plastic than SMR (Norin & Metcalfe, 2019), but on the other hand, SMR and MMR are thought to be positively correlated (Killen, Glazier, et al., 2016; Norin & Clark, 2016). We nonetheless expected to see changes in AS due to predicted changes in SMR.

## Materials and Methods

### Experimental animals

Juvenile Eurasian minnows (*Phoxinus phoxinus* Linnaeus) were captured in spring 2018 from River Kelvin (55.86667, −4.31667; Glasgow, United Kingdom) using dip-nets. The sampling location was an artificial side channel along the River Kelvin where small minnows are trapped as they pass over a weir and are unable to return to the main river. Fish were transported to the nearby University of Glasgow aquarium facilities and held at 15 °C in two large stock tanks (100 x 40 x 30 cm) each filled with 100-150 individuals (density = 833 to 1250 fish m^−3^) for 11 months before the study, which took place in April and May 2019. During this holding period, fish were fed *ad libitum* a combination of pellets and blood worms and were on a 12 h light: 12 h dark photoperiod.

### Experimental design

Experiments were conducted on a total of 80 fish. Since the capacity of the respirometry set-up was of 16 fish (each such group is hereafter referred to as a “lot”), five lots were subjected to respirometry before and after exposure to the social treatments (combination of group size and shelter availability). Each experiment consisted of an initial respirometry trial, a 3-week holding in a social treatment, and a final respirometry trial (Fig. 1). A group of 16 minnows were haphazardly picked from the two stock tanks 48 hours before the onset of an experiment, isolated in a rearing tank (40 x 40 x 30 cm), and fasted.

**Figure 1:**
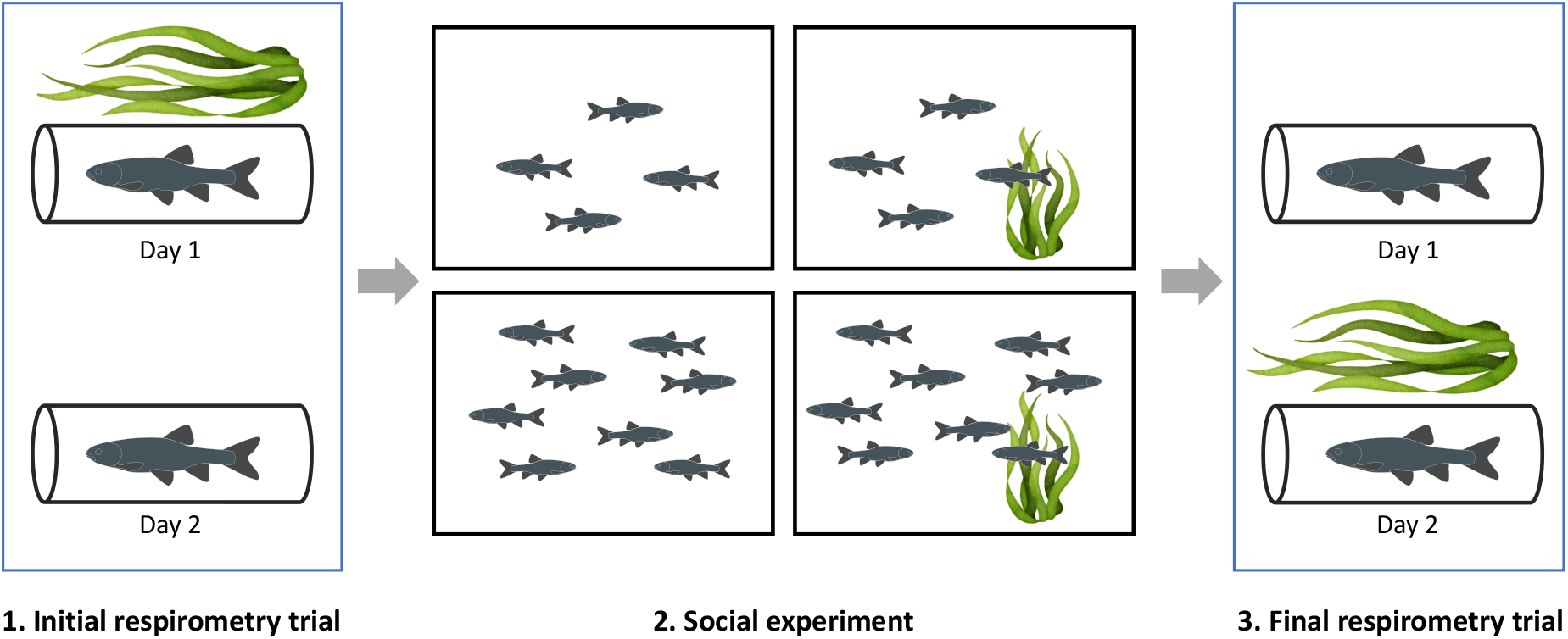
Experimental design of the study. Each experiment consisted of an initial respirometry trial, a 3-week holding in a social treatment, and a final respirometry trial. 1. Initial respirometry trial: Fish oxygen uptake was measured for ~45h during which chambers were covered with artificial plant shelter for approximately half of the trial duration. 2. Social experiment: After the initial respirometry trial, fish were allotted in groups of four or eight fish and placed in experimental tanks containing artificial plant shelter or not, thus forming different social treatments. Fish stayed in their social treatment for 3 weeks. 3. Final respirometry trial: After the social treatment, fish oxygen uptake was measured again by respirometry, in chambers covered with artificial plant shelter for half of the trial duration. Each experiment involved 16 fish (maximum capacity of the respirometry set-up), thus this process was repeated five times, for a total of 80 fish.

Each respirometry trial was conducted to estimate fish metabolic rates in the presence or absence of artificial plant shelter. Fish were placed in individual glass chambers (~100 ml) separated by opaque white dividers to prevent fish from seeing each other. Respirometry trials lasted ~45h during which chambers were covered with artificial plant shelter for approximately half of the trial duration (Fig. S1). At the end of the initial respirometry trial, fish were weighed, measured and injected with a unique combination of visible implant elastomer (Northwest Marine Technology, Anacortes, WA, USA) in the dorsal body surface to allow individual identification. The 16 fish within a given lot were afterwards allotted in groups of four or eight fish and placed in experimental tanks (40 x 40 x 30 cm) containing artificial plant shelter or not, thus forming different social treatments. After the three week holding in their social treatment, the 16 fish were weighted and measured again, and the final respirometry trial was conducted. The whole experiment, from the beginning of the initial respirometry trial with the first lot to the end of the final respirometry trial with the last lot, lasted 41 days.

In total, there were 14 experimental holding tanks. In eight of these experimental tanks, the social treatment was defined by a group size of four fish (density = 83 fish m^−3^) either with, or without, artificial plant shelter (four experimental tanks each). In the remaining six experimental tanks, the social treatment was defined by a group size of eight fish (density = 166 fish m^−3^) either with, or without, artificial plant shelter (three experimental tanks each).

Fish were fed *ad libitum* a combination of pellets and blood worms in their experimental holding tank during the 3-week social experiment to minimize potential effects of density on individual food intake and growth. Daily specific growth rate (SGR: in % day^−1^) during the 3-week social experiment was calculated for each individual using the following equation:

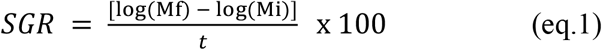

where *M_f_* is the observed mass at the time of the final respirometry trial, *M_i_* is the observed mass at the time of the initial respirometry trial, and *t* is the number of growth days. Over the 3-week social experiment, SGR was higher for fish held in groups of four (mean ± standard deviation: 0.64 ± 0.27 % day^−1^, from −0.07 to 0.99% day^−1^, Fig. S2) than for fish held in groups of eight (0.50 ± 0.19 % day^−1^, from 0.09 to 0.99% day^−1^), and this difference was significant (p=0.004, R^2^_adj_ = 0.084). No relationship was found between SGR and metabolic rates measured at the final experiment (see Supplementary Information for details: Tables S1-S2, Fig. S2-S3).

### Respirometry trials

Metabolic rates were estimated using oxygen consumption rates (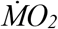: mg O_2_ hr^−1^; Svendsen et al., 2016), determined via intermittent flow-through respirometry equipment and software (Firesting, PyroScience, Aachen, Germany). Water was continuously circulated through each chamber with a peristaltic pump and gas impermeable tubing. Automated flush pumps refreshed the chambers with UV-treated and oxygenated water for 2 min of every 7-min cycle. Dissolved oxygen concentrations were maintained above 80% air saturation at all times with air-bubblers. Temperature was measured with a Pt100 temperature probe and maintained at 15 °C with a TMP-REG instrument (Loligo Systems, Tjele, Denmark) by recirculation of water through a stainless coil in a cold bath.

Respirometry trials lasted ~45h (43.8 to 46.1h), and chambers were covered with artificial plants for about half of its duration (~ 21.5 ± 2 hours; Fig. 2). Presence of artificial plant shelter was randomly set to occur during the first or the second half of the initial respirometry trial, and order was reversed for the final respirometry trial. Respirometry trials started mid-afternoon, and condition (with or without artificial plant shelter) was changed at around noon the next day (~21h after the onset of the respirometry trial). Approximately 43h after the onset of the respirometry trial, fish were taken out of their chamber one by one for a 2-min chase protocol (Roche et al., 2013) and returned in their chamber for immediate measurement of 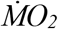 to estimate their maximum metabolic rate MMR (Fig. 2). Respirometry resumed for another hour, and fish were removed from the chambers and transferred to their original experimental tank. Background oxygen consumption in each empty chamber was recorded over three 7-min cycles at the start and end of each respirometry trial.

**Figure 2:**
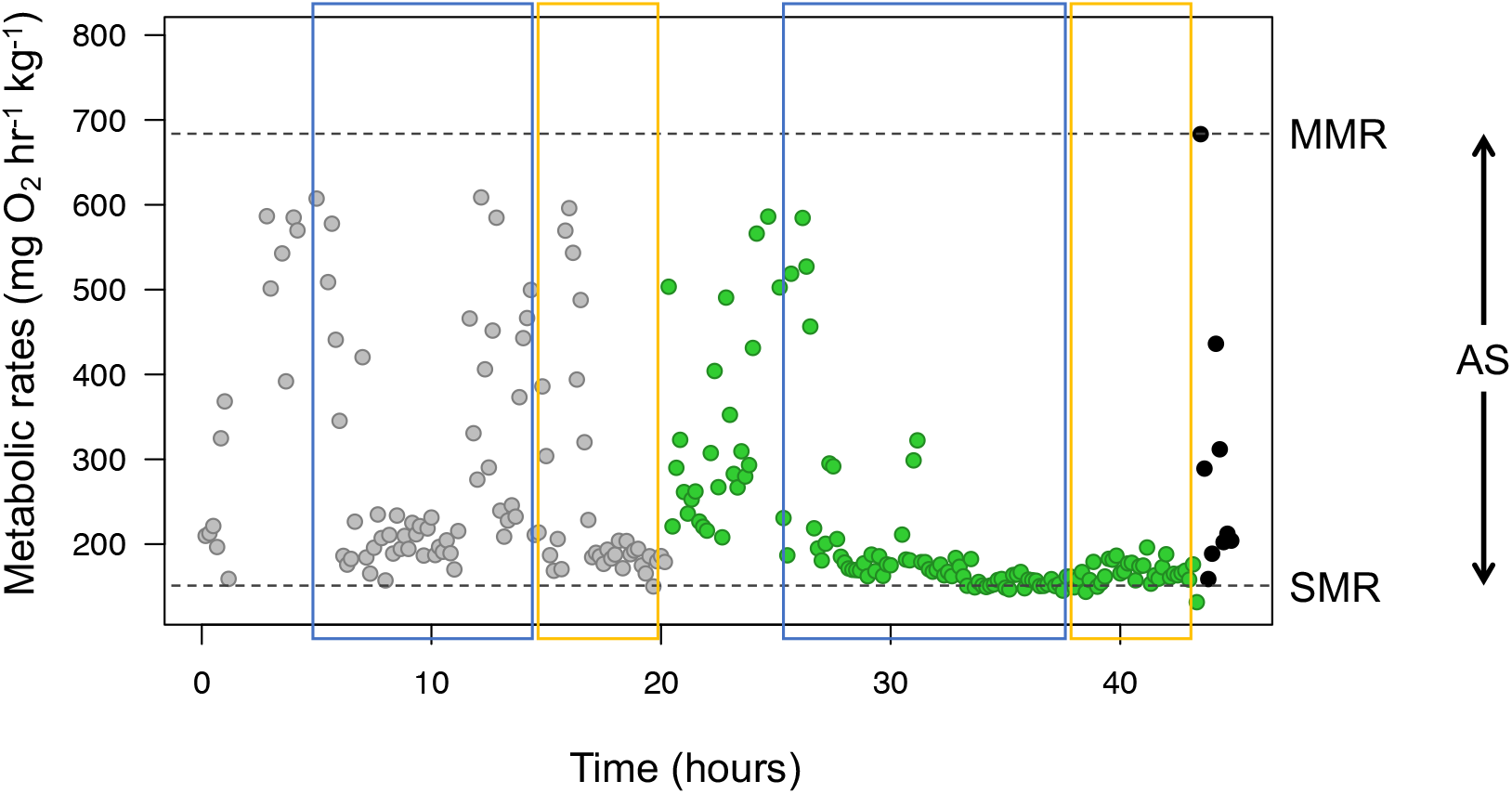
Experimental protocol to obtain 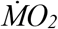 data for the Eurasian minnow. The example shows a 48-h long respirometry trial which started with the condition “without plant” (grey points). The condition was changed to “with plant” (green points) the next day at around noon. On the last day at noon, fish was removed from the respirometry chamber, chased, and immediately placed back into the chamber to obtain MMR (black points). Blue and yellow rectangles represent the range of data used for estimation of night-time and day-time minimum 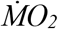, respectively, with or without plant cover. Top and bottom horizontal dotted lines show MMR and SMR.

### Calculation of metabolic rates

Metabolic rates were calculated by multiplying the slopes of decline in oxygen concentration in the chamber during closed measurement cycles, excluding the first 30 seconds, by the volume of the chamber (corrected for the volume of fish, assuming a density of 1 kg l^−1^) using the package *FishResp* in R (Morozov et al., 2019; R Foundation for Statistical Computing, 2018). Background oxygen consumption was subtracted from 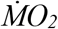 measurements, assuming a linear change between measures taken at the start and end of each trial. Day-time and night-time minimum metabolic rates (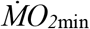; mg O_2_ kg^−1^ hr^−1^) were calculated separately to account for the potentially different effect of the presence of shelter during day-time and night-time. 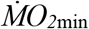 were estimated using the 0.2 quantile of the 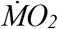 data with the package *fishMO_2_* in R (Chabot et al., 2016; Chabot, 2016). The range of data used for the calculation of night-time 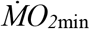 started 5 hours after fish were put in the chamber (at around 9:30 pm) or 5 hours after the change in condition (presence of plant shelter or not; at around 6:30 pm), and ended in the morning at 7:00 am, moment at which lights were turned on. The range of data used for the calculation of day-time 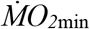 started at 7:00 am and ended at the change in condition, or when fish were retrieved from the chamber for the chase protocol (Fig. 2). Standard metabolic rate (SMR; mg O_2_ kg^−1^ hr^−1^) was set as the lowest estimate of day-time or night-time 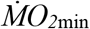 over a trial. MMR (mg O_2_ kg^−1^ hr^−1^) was estimated as the highest rate of oxygen consumption over 3 a minute rolling average regression within a measurement cycle following the chase protocol. Aerobic scope (AS; mg O_2_ kg^−1^ hr^−1^) was calculated as the difference between MMR and SMR. All metabolic rates were adjusted to the mean body mass of the fish in our sample (mean ± s.d.: 1.95 ± 0.57 g) using the slope *b* of the log-log relationship between 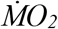 and mass (Steffensen et al., 1994; Ultsch, 1995).

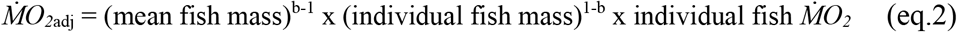

### Statistical analyses

All data are available from Zenodo (https://doi.org/10.5281/zenodo.4705121, Chrétien et al., 2021). All analyses were computed in R v. 3. 6. 0 (R Foundation for Statistical Computing, 2018). Effects of presence of shelter on night-time and day-time 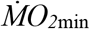 measured during initial and final respirometry trials were tested using linear mixed effects models (LMM) with the package *lme4* (Bates et al., 2014). Full models included trial (initial or final), trial day (1^st^ or 2^nd^), presence or absence of plant shelter during the trial, fish body mass (g), and all interactions as fixed effects. There was no relationship between fish body mass (g) and mass-adjusted metabolic rates, so fish body mass was excluded from models. Models included fish ID, lot number (1 to 5), and tank (referring to the experimental tank in which fish were held during the social treatment) as potential random effects. The best random structure was first selected by comparison of Akaike information criterion on full models (AIC; Zuur et al., 2009), then the fixed structure was simplified by removal of non-significant interactions. Final models included fish ID and lot number as random effects in a nested structure (lot number/fish ID). Model assumptions were met when response variables were log-transformed. For all models, assumptions of homoscedasticity, linearity and normality were confirmed by visual inspection of residual plots.

Effects of group size and shelter availability on SMR, MMR, and AS were tested with LMM using data from the initial and final respirometry trials, social treatment conditions (group size: four or eight fish, shelter availability: presence or absence of artificial plant in experimental tank), fish body mass, and all interactions as fixed effects. Full models included fish ID, lot number, and tank as potential random effects, and best random structure was selected by comparison of AIC. Final SMR model included fish ID and lot number as random effects in a nested structure (lot number/fish ID). Only fish ID was retained as random effect in final MMR and AS models. Model assumptions were confirmed by visual inspection of residual plots.

Effect sizes were calculated using estimated marginal means from models obtained with the package *emmeans* (Lenth & Hervé, 2015). Marginal R^2^ (R^2^_m_: variance explained by fixed effects) and conditional R^2^ (R^2^_c_: variance explained by fixed and random effects) were calculated from the models fitted through restricted maximum likelihood analysis (Bolker et al., 2009; Harrison et al., 2018). The difference between R^2^_c_ and R^2^_m_ for each model represent variability due to the random effects (Nakagawa & Schielzeth, 2013).

## Results

### Presence of shelter and metabolic rates

Respirometry timing (initial or final), trial day, and plant shelter (presence or absence during respirometry) had significant effects on night-time 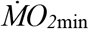 (p=0.002, p<0.001, and p=0.002, respectively; Table 1). Night-time 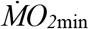 recordings were on average 8.7% higher during the final respirometry trial, 16.2% lower on the 2^nd^ day of trial, and 7.9% lower in the presence of plant shelter (Fig. 3A-B). Day-time 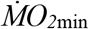 was influenced by respirometry timing (p<0.001; Table 1) and trial day (p=0.006) but not by the presence of plant shelter (p=0.819). Day-time 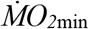 was on average 26.9% higher at the final respirometry trial, and 5.8% lower on the 2^nd^ day of trials (Fig. 3C-D). There was an interaction between trial day and plant shelter on day-time 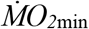 (p=0.044): in the presence of plant shelter, day-time 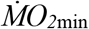 rates measured on the 2^nd^ day were 10.0% lower than that those of the 1^st^ day.

**Table 1:** Results of linear mixed models relating night-time and day-time minimum metabolic rates 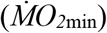 of Eurasian minnows to respirometry trial (initial or final), trial day, and presence or absence of plant shelter. R^2^_m_ is the marginal R^2^ (variance explained by the fixed effects) and R^2^_c_ is the conditional R^2^ (total variance explained by the fixed and the random effects).

| Response variable | Effect | $\chi^2$ | p-value | $R^2_m$ | $R^2_c$ |
| --- | --- | --- | --- | --- | --- |
| log Night-time $\dot{M}O_{2min}$ | Trial | 9.313 | 0.002 | 11.5 | 42.6 |
|  | Day | 42.501 | <0.001 |  |  |
|  | Plant shelter | 9.229 | 0.002 |  |  |
| log Day-time $\dot{M}O_{2min}$ | Trial | 111.905 | <0.001 | 16.7 | 56.2 |
|  | Day | 7.591 | 0.006 |  |  |
|  | Plant shelter | 0.052 | 0.819 |  |  |
|  | Day* Plant shelter | 4.051 | 0.044 |  |  |

**Figure 3:**
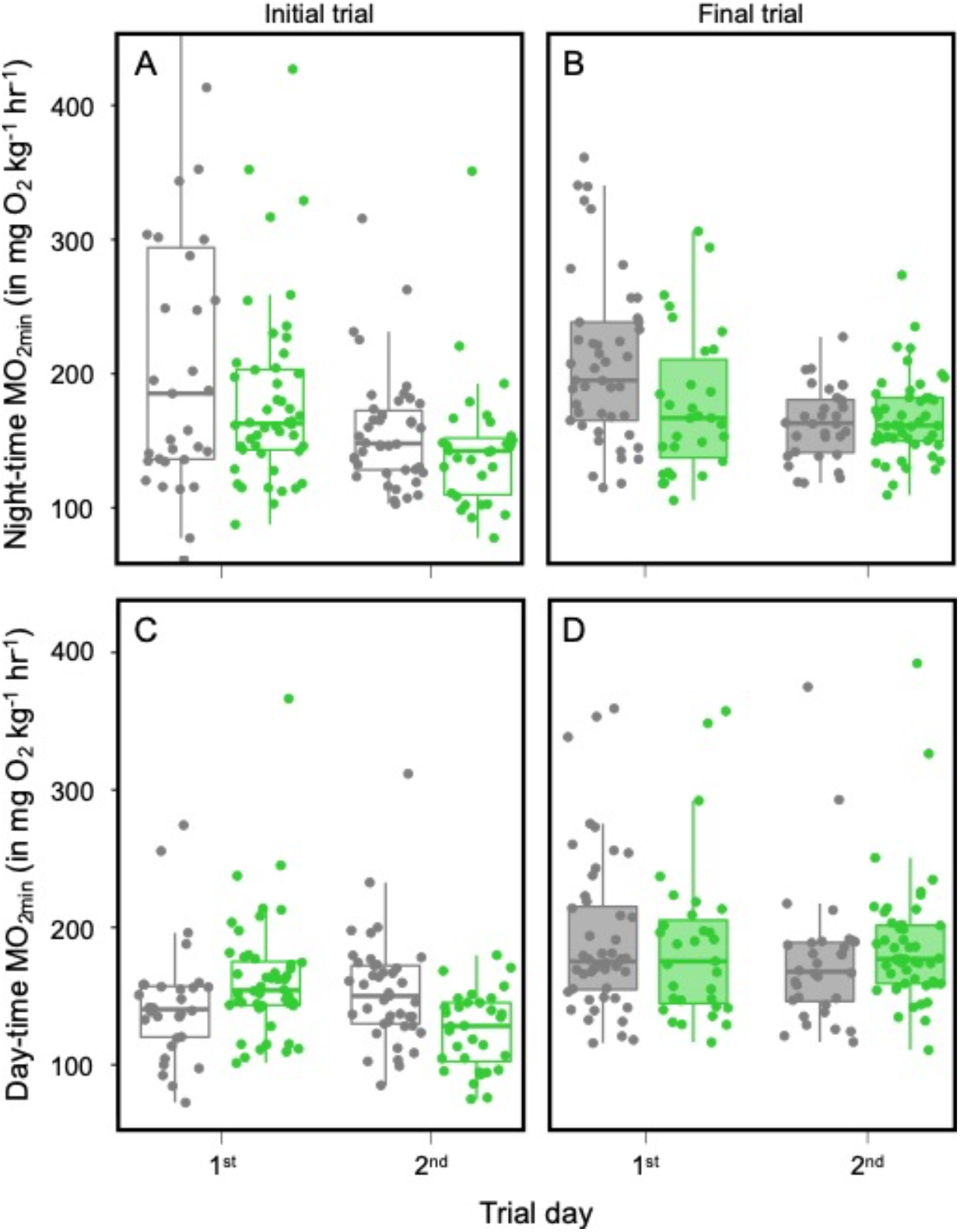
Observed night-time (A, B) and day-time (C, D) metabolic rates in initial (clear) and final (shaded) respirometry trials. Grey and green dots represent estimates in absence or in presence of plant shelter, respectively. Middle thick line of the boxplots corresponds to the median, lower and upper limits correspond to the first and third quartiles of the data, and whiskers extend to the range of the data.

### Social environment and metabolic rates

There was an overall increase in SMR after the 3-week social treatment (p<0.001; Table 2), and an interacting effect of trial and group size (p=0.006). SMR estimates were 28% higher at the final respirometry trial compared to the initial one for fish held in groups of four, while SMR increased of 13% between the two trials for fish held in groups of eight (Fig. 4A-B). Plant shelter availability in experimental tanks did not influence SMR. MMR did not change between the initial and final respirometry trials (p=0.254). Fish held in groups of four had, however, higher MMR than fish held in groups of eight (p=0.005; Fig. 4C-D). Finally, there was an overall reduction in AS after the 3-week social treatment (p=0.029; Table 2). Group size also negatively influenced AS (p=0.008; Fig. 4E-F). Plant shelter availability in experiment tanks did not influence MMR or AS (Table 2).

**Table 2:**
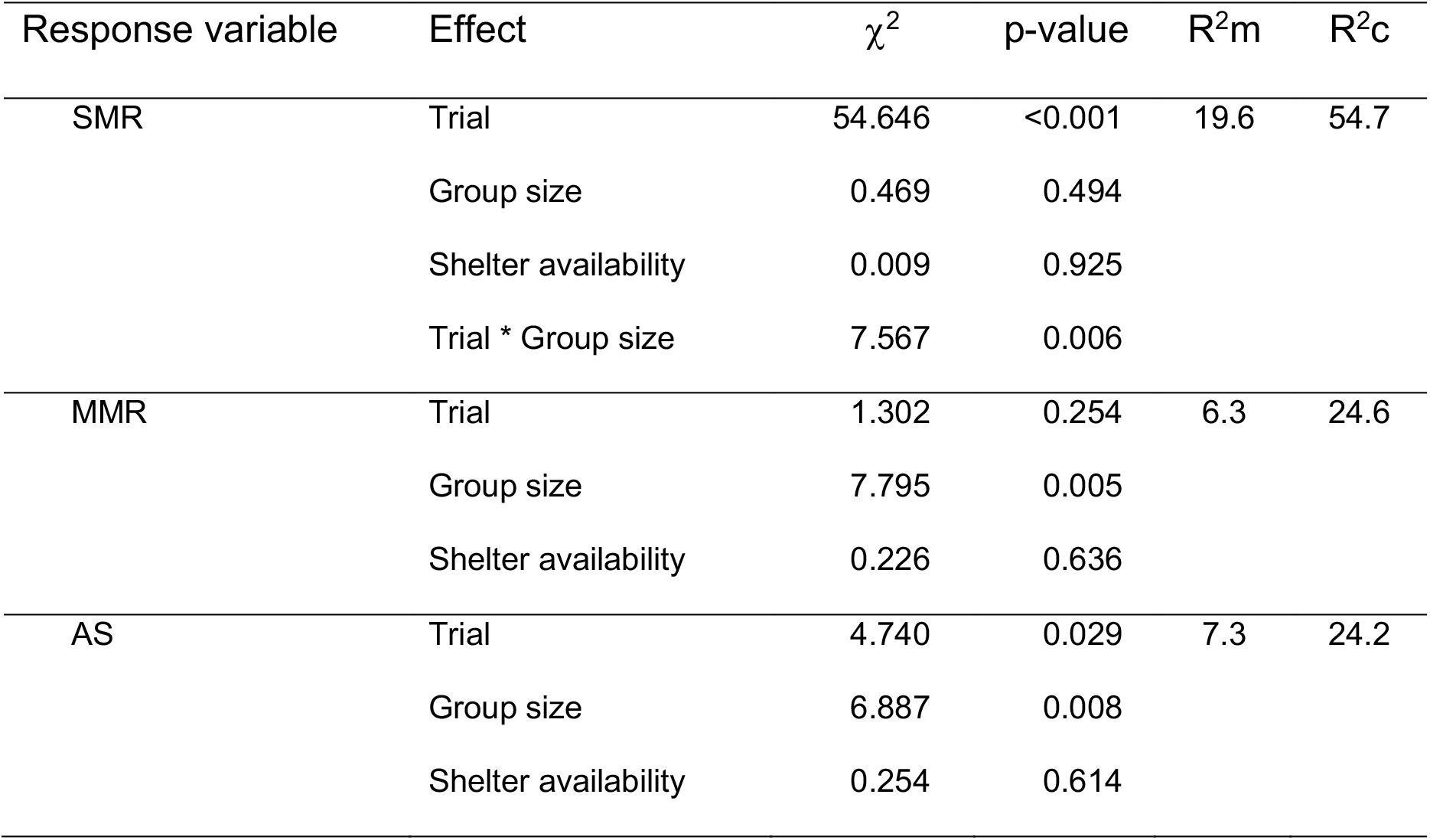
Results of linear mixed model relating metabolic rates of Eurasian minnows to the moment of the respirometry trials, and social treatment (group size and shelter availability). Fish ID and lot number were included in the SMR model as random effects. Only fish ID was included as a random effect for MMR and AS models. R^2^_m_ is the marginal R^2^ (variance explained by the fixed effects) and R^2^_c_ is the conditional R^2^ (total variance explained by the fixed and the random effects).

**Figure 4:**
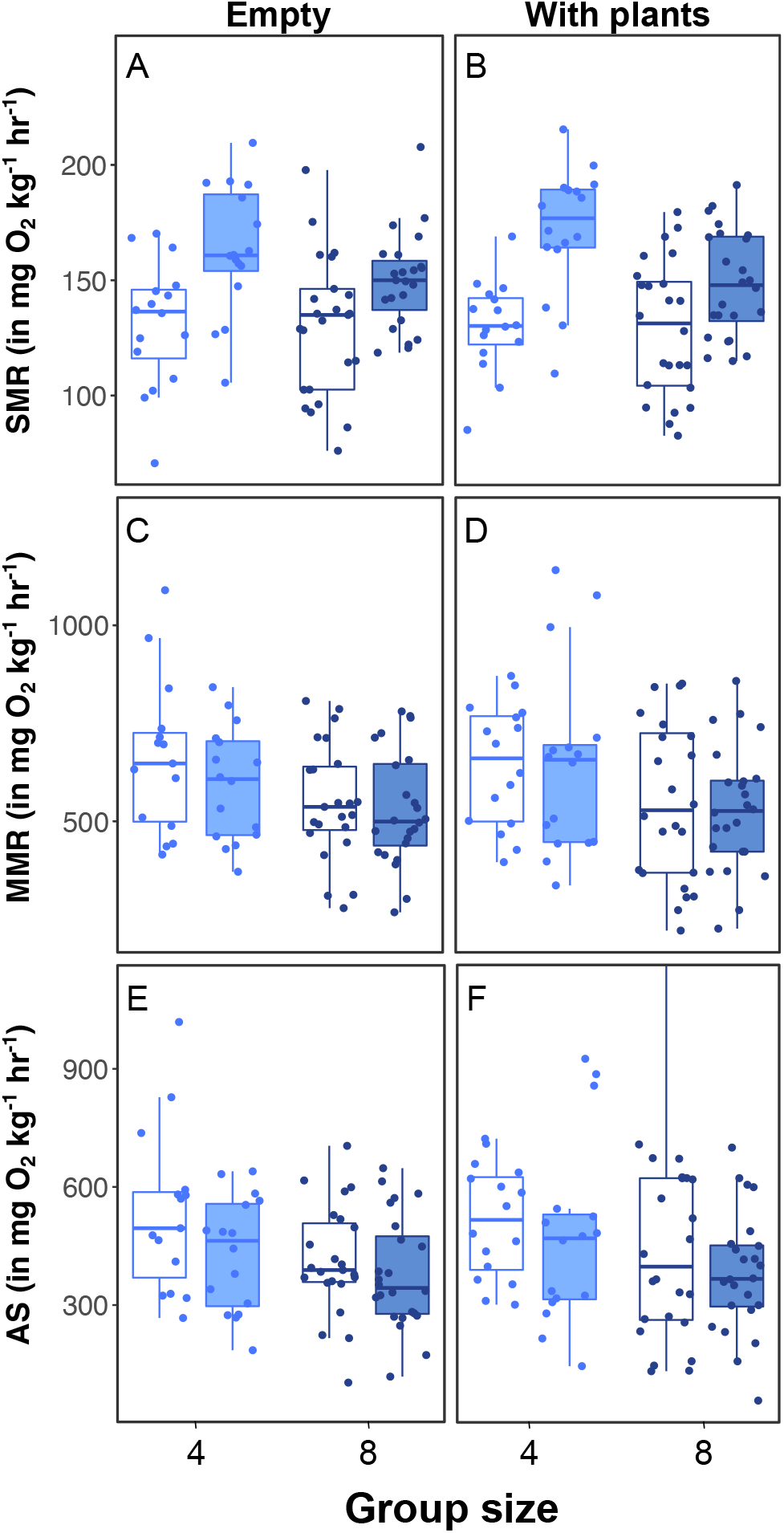
Observed SMR (A-B), MMR (C-D), and AS (E-F) of Eurasian minnow. Light blue and dark blue boxes and points represent estimates for fish held in groups of four and eight, respectively. Clear and shaded boxes represent initial and final respirometry trials, respectively. A-C-E panels refer to tanks without plant shelter, and B-D-F refer to tanks containing plant shelter. Middle thick line of the boxplots corresponds to the median, lower and upper limits correspond to the first and third quartiles of the data, and whiskers extend to the range of the data.

## Discussion

The main goal of this study was to assess whether exposure to a given social group size and level of shelter availability had the potential to modulate expression of metabolic traits. Both before and after holding in different social treatments, minimum metabolic rates measured in presence of shelter were lower than those measured in absence of shelter. Presence of plant shelter during respirometry trials reduced metabolic rates regardless of the social group size and shelter availability fish were exposed to. We did, however, observe an overall increase in the SMR of Eurasian minnows between the initial and final respirometry trial, with the increase in SMR throughout the study being two-fold higher for fish held in groups of four as compared to that of fish held in groups of eight. Availability of shelter in holding tanks during the social treatments did not affect metabolic rates. Our results suggest that group size has metabolic costs that carry over, even when fish are at rest and in isolation, such as during respirometry trials. This means that group size can have a modulating effect on levels of baseline metabolism, which could in turn have implications on an animal’s energy budget, including growth, reproductive investment, and overall performance capacity. In the current study, the presence of more groupmates was associated with lower metabolic rate, suggesting that a reduction in energy demand may be an additional benefit of living in larger social groups.

### Presence of shelter and metabolic rates

Presence of plant shelter during respirometry lowered estimates of metabolic rates both before and after exposure to the social treatments. Presence of shelter has been associated with lower metabolic rates in some species (Finstad et al., 2004; Fischer, 2000; Millidine et al., 2006; Norin et al., 2018) but not in others (Fischer, 2000; Kegler et al., 2013), or to mixed results (Chrétien et al. 2020). Using shelter can reduce the occurrence of otherwise energetically demanding activities, such as those associated with maintaining vigilance against predators (Lind & Cresswell, 2005; Millidine et al., 2006). It was surprising that the effect of shelter was stronger for night-time than for day-time 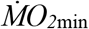, assuming the main reason for sheltering is to remain visually hidden. This pattern was nonetheless observed in another study, where an effect of shelter presence was observed during the night but not during the day (Norin et al., 2018). It is possible that fish showed higher levels of spontaneous activity during day-time which might mask any effect of the shelter on 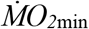, although no consistent relationship has been observed between activity and light intensity in our study species (Jones, 1956). Another potential explanation is that fish had time to acclimate to the presence of shelter before night-time, and therefore had expected that they could be sheltered at night. We predicted that the magnitude of the effect of shelter on metabolic rates would be smaller after the 3-week social experiment. This trend was not observed, suggesting that individuals did not adjust their metabolic response to immediate shelter presence, regardless of the group size or level of shelter availability they received during the experiment. This indicates that shelter availability has a consistent and robust lowering effect on resting metabolic rates in Eurasian minnow and likely other species with similar social systems and patterns of habitat use.

### Social environment and metabolic rates

There was an overall increase in estimates of SMR throughout the study, after fish had been exposed to the social treatments. Importantly, group size affected the strength of the increase: fish held in groups of four showed a two-fold higher increase in estimated SMR than fish held in groups of eight. We cannot rule out that conditions may have been more favorable for growth in tanks with groups of four, even if food was not a limited resource in any social treatment. However, there was no relationship between final SMR and SGR, nor was there an interaction between SGR and social treatment conditions (Tables S1-S2, Fig. S2-S3), suggesting other mechanisms are more likely to explain the differences observed. For instance, fish in groups of four potentially had more volume available for individual exploration and an increased need for individual vigilance, potentially increasing the cognitive load and associated metabolic costs that may carry over, even when the fish are at rest, during respirometry for estimates of SMR (Moss et al., 1998). Prolonged changes in locomotor activity level due to social interaction or vigilance may induce changes in muscle enzyme levels and mitochondria density, and thus affect fish minimum energy demand (Killen, Glazier, et al., 2016). Intensity of competition and strength of hierarchy structures could also vary differently with group sizes. With increasing group size, competition for limited resources like shelter may increase but dominance hierarchies tend to weaken, as the cost of interacting with multiple individuals may become too high (Sloman & Armstrong, 2002). For example, Pottinger and Pickering (1992) observed that social hierarchies emerged in rainbow trout *Oncorhynchus mykiss* held for six weeks in pairs or in groups of 5, but not in groups of 10 fish. An increase in aggressive behaviour such as pecking incurs increased activity and metabolic costs (Marchand & Boisclair, 1998). Presence of plant shelter in experimental tanks did not affect SMR. This was surprising given that tank enhancements such as artificial plants can be used as tools to reduce aggression and provide shelter in captivity (Näslund & Johnsson, 2016). It is possible that plant shelter in the experimental tanks were rather considered as a limited resource to compete for, which could have enhanced social stress. Sustained stress in social groups with stronger dominance hierarchies could carry over and limit our ability to effectively estimate SMR (Killen et al., 2014; Metcalfe et al., 2016; Sloman et al., 2000). Additional research on the effects of social dynamics on fish cognitive abilities or stress indicators could shed light on the mechanisms underlying the results observed here.

Fish held in groups of four had significantly higher MMR and AS than fish held in group of eight before the 3-week holding in their social treatment (Table 2). We did not expect group size to affect metabolic rates in the initial respirometry trial as fish were all held in the same high-density stock tank beforehand. It therefore appears that this result was driven by a single lot of fish. The first lot of 16 fish subjected to our experiment reached overall higher MMR (and AS) than the other lots at the initial respirometry trial (Fig. S4), and were all allotted in groups of four. We included “lot number” as a potential random effect in all our models to account for higher similarities in fish from the same lot compared to other fish. Lot number was retained in a nested structure with fish ID for night-time 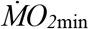, day-time 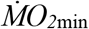 and SMR models. It was not, however, kept in models on MMR or AS, because its inclusion resulted in singular fits (Matuschek et al., 2017): no variance was associated to the random effect “lot number”. In any case, models using either “fish ID” or “lot number /fishID” as a random component generated similar results (Table S3). High susceptibility to capture is a trait that can correlate with MMR (Redpath et al., 2010), and might explain the pattern we observed when comparing MMR of the first lot of fish captured to MMR of the subsequent ones. While this pattern could be interesting to investigate in other studies, we can only interpret it here as a measurement artefact and cannot link this result to the social treatments.

There is evidence that shoaling can have a “calming effect” and reduce metabolic rates of social fish species, through conspecific visual and olfactory cues (Nadler et al., 2016). The social treatment revealed that group size could influence SMR, which can be attributable to increased social stress at lower densities for these social fish. It is possible that increased group size and habitat complexity induces metabolic plasticity, which suggests that selection on energy expenditure in animals with strong social systems may be less likely to result in genetic change. Our results highlight the importance of understanding the role of social dynamics on variations in individual metabolic traits and thus on the physiological consequences of habitat conditions.

## Acknowledgements

We would like to thank the fish care team at the Graham Kerr building facilities for their help in maintaining fish throughout the study. The procedures described in this paper comply with animal care guidelines approved within the UK and were carried out under Home Office Project License no. PB948DAA0. This work was supported by grants from Natural Sciences and Engineering Research Council of Canada (NSERC, CGS-D to E.C. and Discovery Grant to D.B.), Fonds de recherche du Québec – Nature et Technologies (FRQNT, to E.C.), and Groupe de recherche en limnologie et environnement aquatique (GRIL, to E.C.). SSK was supported by a Starting Grant from the European Research Council starting (grant 640004) and a Natural Environment Research Council Advanced Fellowship (NE/J019100/1).

## Author’s contributions

E.C. and S. S. K. designed the experiments. E.C. performed the experiments, analyzed the data and led the writing of the manuscript. All authors contributed to the interpretation of data, the critical revision of the drafts and gave final approval for publication.

## Data availability statement

The datasets generated and analyzed during the current study are available from Zenodo repository https://doi.org/10.5281/zenodo.4705121 (Chrétien et al., 2021).

