## Supplementary materials for "Group size influences individual metabolic traits in a social fish"

Supporting Information

### Respirometry trial set-up:

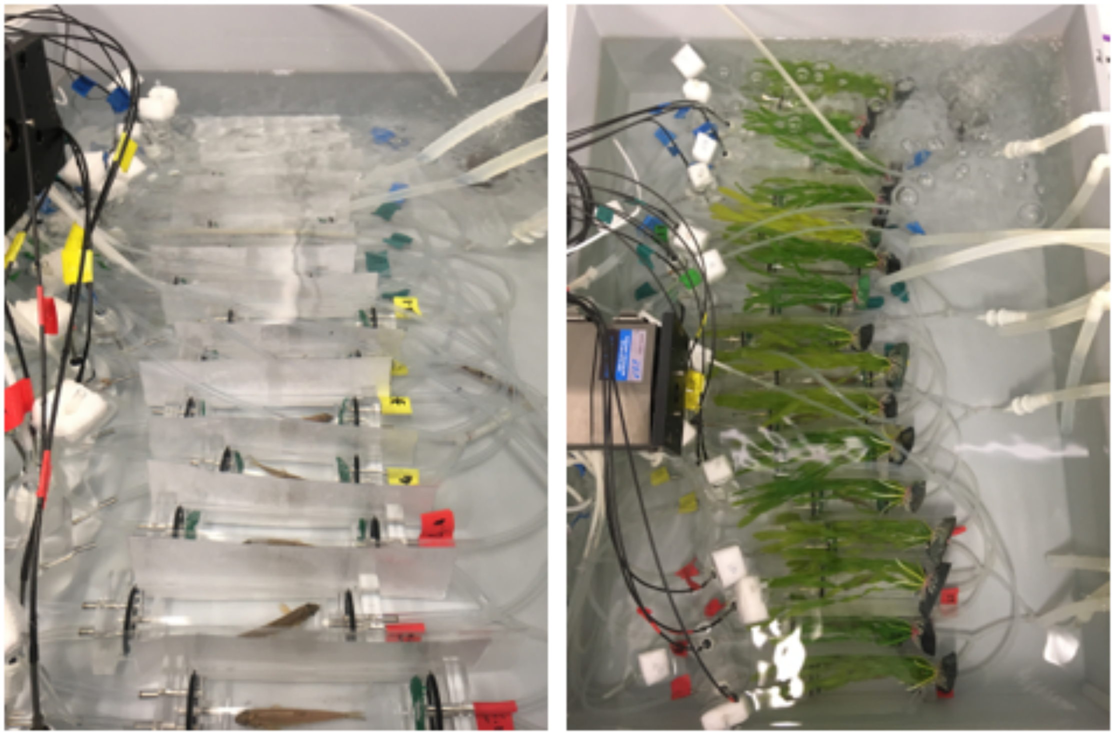

Figure S1: Respirometry trial setup. 16 fish tested each time. There were dividers between chambers so fish could not see each other. Trials lasted two days, one day during which chambers were not covered with plants, while the other day the chambers were covered with plants. Plant cover was randomly set to occur on the 1$^{st}$ or 2$^{nd}$ day of trial for each lot of fish tested together, and order was reversed for the final respirometry trial.

**Specific growth rate**

Fish were fed *ad libitum* a combination of pellets and blood worms in their experimental holding tank during the 3-week social experiment to minimize potential effects of density on individual food intake and growth. We compared daily specific growth rate (SGR: in % $day^{-1}$) across social treatments and observed that fish held in groups of eight had on average lower SGR (Table S1), but that SGR was overall very variable among individuals (Figure S2).

Further, we tested if SGR could influence metabolic rates. It was not possible to include SGR in the models presented in the results of the manuscript as those models included two estimates of metabolic rates per individual (initial vs final respirometry trial), while we have a single SGR per individual. Therefore, we tested if SGR and its interaction with the social treatment could influence metabolic rates estimated at the final respirometry trial. There was no effect of SGR on any metabolic rate (Table S2, Figure S3). In addition, there was no interaction between SGR and group size or shelter availability.

Table S1: Results of linear model relating specific growth rate (SGR) of Eurasian minnows to the social treatment (group size and shelter availability).

| Response variable | Effect | Estimate ± standard error | *F* | p-value | $R^2_{adj}$ |
|---|---|---|---|---|---|
| SGR | Group size | -0.152 ± 0.051 | 9.011 | 0.004 | 8.4 |
| | Shelter availability | 0.025 ± 0.050 | 0.248 | 0.620 | |

Table S2: Results of linear model relating final standard metabolic rate (SMR), maximum metabolic rate (MMR) and aerobic scope (AS) of Eurasian minnows to the specific growth rate (SGR), the social treatment (group size and shelter availability), and their interaction*.

| Response variable | Effect | *F* | p-value |
|---|---|---|---|
| Final SMR | SGR | 2.036 | 0.158 |
| | Group size | 12.232 | <0.001 |
| | Shelter availability | 0.190 | 0.664 |
| Final MMR | SGR | 0.028 | 0.868 |
| | Group size | 4.983 | 0.029 |
| | Shelter availability | 0.285 | 0.595 |
| Final AS | SGR | 0.001 | 0.972 |
| | Group size | 3.182 | 0.078 |
| | Shelter availability | 0.235 | 0.629 |

*All interactions were non-significant and were therefore removed from models.

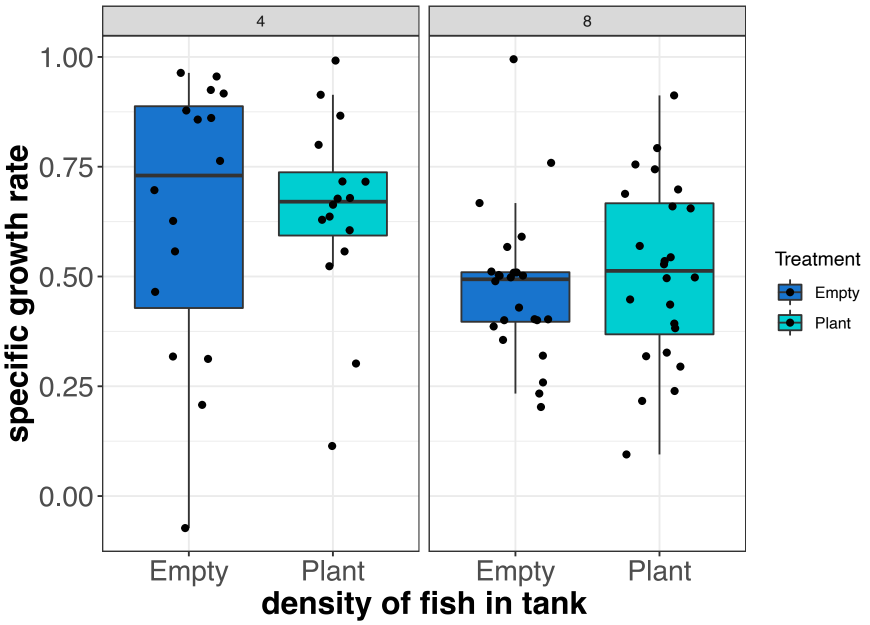


Figure S2: Boxplot of specific growth rate (SGR; in % day$^{-1}$) of fish in each social treatment consisting in a combination of group size (left: four fish; right: eight fish) and shelter availability (no plant shelter in blue; with plant shelter in turquoise). Middle thick line of the boxplots corresponds to the median, lower and upper hinges correspond to the first and third quartiles of the data, and whiskers extend to the range of the data. Each fish SGR is overlayed on the boxplot (black dots).

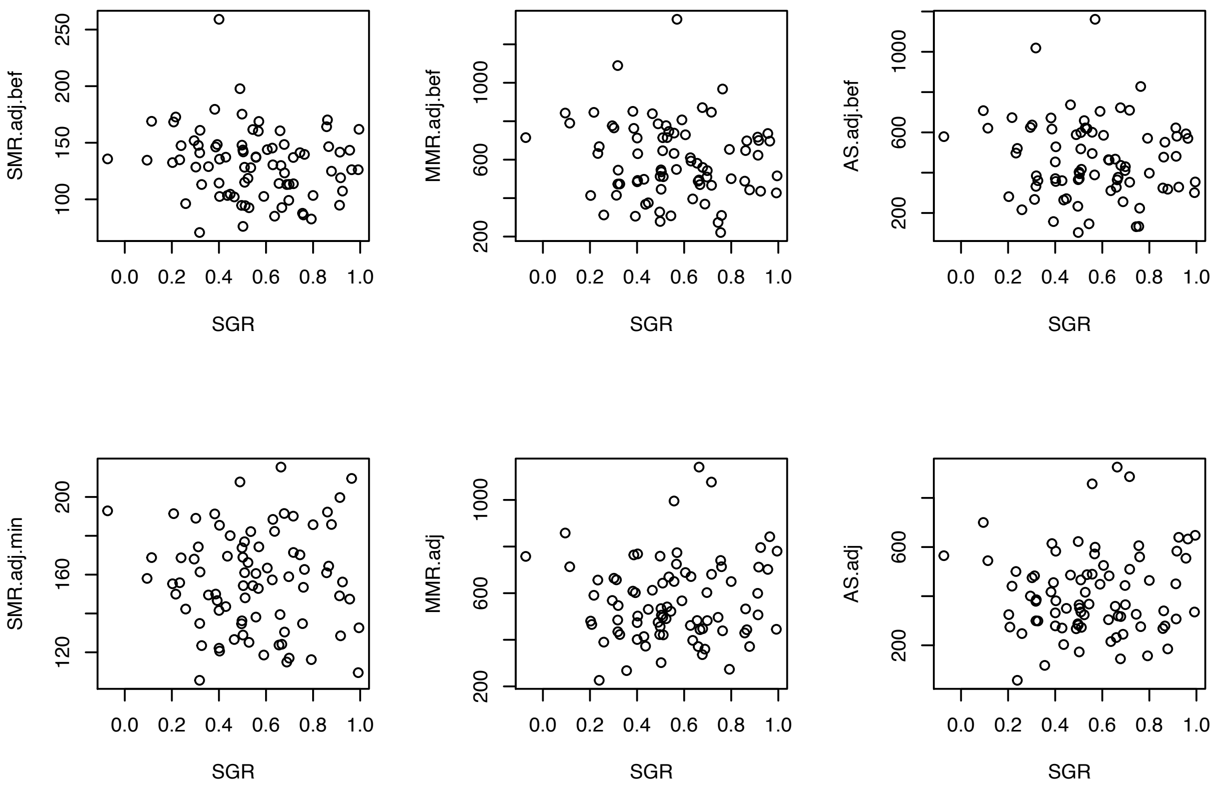


Figure S3: Observed SMR, MMR, and AS of Eurasian minnow (in mg $O_2$ $kg^{-1}$ $hr^{-1}$) during the initial (top row) and final (bottom row) respirometry trial in relation to specific growth rate (SGR; in % $day^{-1}$).

**MMR and AS**

Table S3: Results of linear mixed model relating maximum metabolic rate (MMR) and aerobic scope (AS) of Eurasian minnows to the moment of the respirometry trials and the social treatment (group size and shelter availability). Fish ID and lot number were included in the models as a random effect in a nested structure (lot number/Fish ID). $R^2_m$ is the marginal $R^2$ (variance explained by the fixed effects) and $R^2_c$ is the conditional $R^2$ (total variance explained by the fixed and the random effects).

| Response variable | Effect | $\chi^2$ | p-value | $R^2$m | $R^2$c |
|---|---|---|---|---|---|
| MMR | Trial | 1.312 | 0.252 | 6.3 | 24.7 |
| | Group size | 6.717 | 0.010 | | |
| | Shelter availability | 0.224 | 0.636 | | |
| AS | Trial | 4.813 | 0.028 | 7.5 | 23.2 |
| | Group size | 7.159 | 0.007 | | |
| | Shelter availability | 0.262 | 0.609 | | |

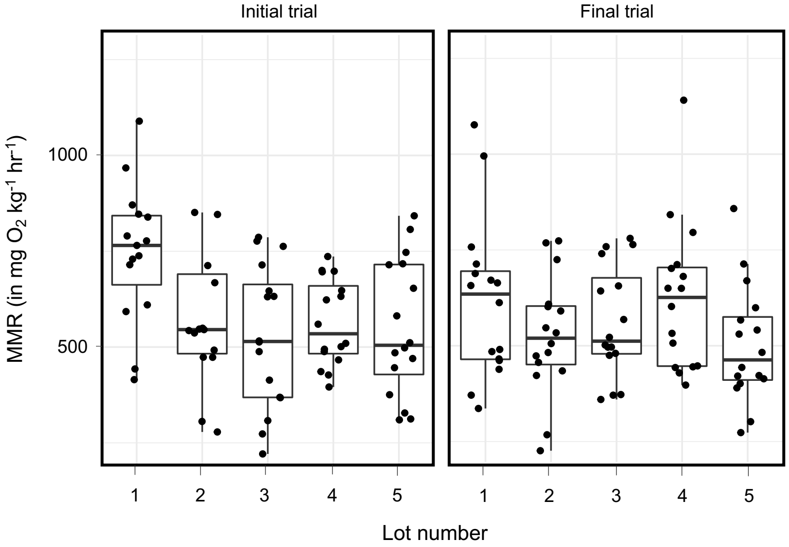


Figure S4: Maximum metabolic rates (MMR) per lot of fish captured together from the stock tank. Left panel shows rates measured in the initial respirometry trial, right panel shows rates measured at the final respirometry trial. Middle thick line of the boxplots corresponds to the median, lower and upper hinges correspond to the first and third quartiles of the data, and whiskers extend to the range of the data. Each fish MMR is overlayed on the boxplot (black dots).
